# Mapping the Glycogen Synthase 1 Interactome in Brown Adipocytes

**DOI:** 10.64898/2025.12.01.691470

**Authors:** Jeslyn Zhang, Stacy Tletlepantzi Lara, Sicheng Zhang, Vanja Panic, Vijaya Pandey, Ezekiel Delgado, Julian Wang, James A Wohlschlegel, Claudio Villanueva

## Abstract

Brown adipose tissue (BAT) is distinguished by its ability to dissipate energy through thermogenesis, a process with considerable therapeutic potential for obesity, type 2 diabetes, and related metabolic disorders. Although glycogen represents a relatively minor energy reserve in adipocytes compared with liver and muscle, emerging evidence shows that glycogen turnover increases sharply during metabolic transitions and is required for full thermogenic activation in brown adipocytes. The rate-limiting enzyme glycogen synthase 1 (GYS1) catalyzes glycogen chain elongation, and adipose-specific Gys1 deletion impairs adaptive thermogenesis, highlighting the importance of GYS1 in coordinating glucose and lipid metabolism; however, the mechanisms regulating GYS1 in BAT remain poorly defined. To address this gap, we used an unbiased proximity-labeling strategy in brown adipocytes by fusing GYS1 to the biotin ligase TurboID and identifying biotinylated proteins through mass spectrometry, revealing 425 putative interactors. These included canonical glycogen-handling proteins such as glycogenin and glycogen phosphorylase, validating the approach, while gene ontology analysis uncovered broader enrichment in cytoskeletal remodeling, vesicle trafficking, and transcriptional regulation, suggesting previously unrecognized roles for GYS1 beyond glycogen synthesis. Notably, we identified several novel candidate interactors, including CAST, PDAP1, and CCDC102A. Together, these findings define the most comprehensive GYS1 interactome reported to date and provide new insights into the molecular mechanisms linking glycogen metabolism to thermogenic function in BAT.

## INTRODUCTION

Brown adipose tissue (BAT) is characterized by its unique ability to expend energy through thermogenesis, a mechanism that carries therapeutic potential for obesity, Type 2 diabetes, and related metabolic disorders (1). Beyond its role in thermogenesis and whole-body energy expenditure, brown adipose tissue (BAT) has emerged as an important modulator of cardiometabolic health. Human imaging studies demonstrate that individuals with detectable BAT exhibit lower prevalence of major cardiovascular risk factors, including dyslipidemia, hypertension, insulin resistance, and coronary artery disease(2). Glycogen metabolism, a primary regulator of energy storage and utilization, is an emerging area of interest in BAT functionality. Glycogen is a branched polymer of glucose functioning as a major mechanism of short-term energy storage and stored primarily in the liver and the skeletal muscle (3).

Despite their low abundance, increasing evidence suggests adipose glycogen stores are acutely regulated to coordinate glucose and lipid metabolism during metabolic transitions (4). Glycogen transiently accumulates following fasting, cold stimulation, or leptin treatment before lipid stores are replenished (5–7). Glycogen degradation has been shown to facilitate lipid droplet biogenesis during brown adipocyte differentiation (8). Recent studies indicate that glycogen turnover increases under thermogenic activation and promotes the expression of thermogenic genes (9,10). However, the molecular mechanisms regulating glycogen metabolism in BAT remain incompletely understood.

Glycogen is synthesized by the combined action of glycogenin, glycogen synthase, and glycogen branching enzyme. These enzymes, together with other glycogen-handling proteins, associate with glycogen to form supramolecular glycogen-protein complexes, termed glycosomes, which serve as hubs for metabolic regulation (11). Glycogenin first acts as a primer to initiate the formation of a short glucose polymer, which is then elongated by glycogen synthase and branched by glycogen branching enzyme (12). Among these enzymes, glycogen synthase (GS) catalyzes the rate-limiting step of the pathway, making it the primary target for regulatory control of glycogen synthesis. GS is a tetrameric glucosyltransferase that catalyzes the addition of UDP-glucose monomers to the growing glycogen chain through the formation of α-1-4 glycosidic linkages (13). GS is allosterically activated by glucose-6-phosphate binding and inactivated by covalent phosphorylation at several N- and C-terminal sites (14,15). There are two isoforms of GS: the widely expressed GYS1 and the liver-specific GYS2. Importantly, *Gys1* deletion in beige adipose tissue impairs its capacity for thermogenic activation in response to cold stimulation, suggesting its role as a key enzyme in coordinating glucose and lipid metabolism (16).

Emerging literature suggests GYS1 may participate in mechanisms beyond its canonical role in cytoplasmic glycogenesis. GYS1 has been characterized as a nucleocytoplasmic shuttling protein, translocating to the cytosol or nucleus in response to intracellular glucose levels (17,18). GYS1 folds into two Rossman-fold domains, structural motifs which enable metabolic enzymes to perform multiple functions, including RNA binding and ribonuclease activity (19,20). Indeed, GYS1 has been reported to associate with mRNA and the translation machinery, pointing to an expanded regulatory network that remains to be fully explored (21,22).

As the complete GYS1 interactome has not yet been characterized in BAT, we adopted an unbiased, proteomics-based approach to explore GYS1 regulatory mechanisms in brown adipocytes. Specifically, we utilized TurboID (TbID)-based proximity labeling to identify BAT-specific interacting partners with GYS1. TbID is an engineered biotin ligase whose enzymatic activity makes it ideal for capturing dynamic interactions that occur between proteins during metabolic transitions (23). In the presence of biotin, TbID uses ATP to generate the reactive intermediate biotin-AMP, which covalently labels endogenous proteins proximal to GYS1 (Figure 1B). Biotin-tagged proteins are isolated by co-immunoprecipitation using streptavidin-coated magnetic beads, then analyzed by LC-MS/MS for identification of GYS1 interacting partners. Pathway analysis revealed expected enrichment in glycogen metabolism, along with unexpected enrichment in transcriptional regulation, suggesting previously unrecognized functions for GYS1. These findings further elucidate the mechanisms that regulate glycogen metabolism in the context of BAT’s unique metabolic functions.

**Figure 1.**
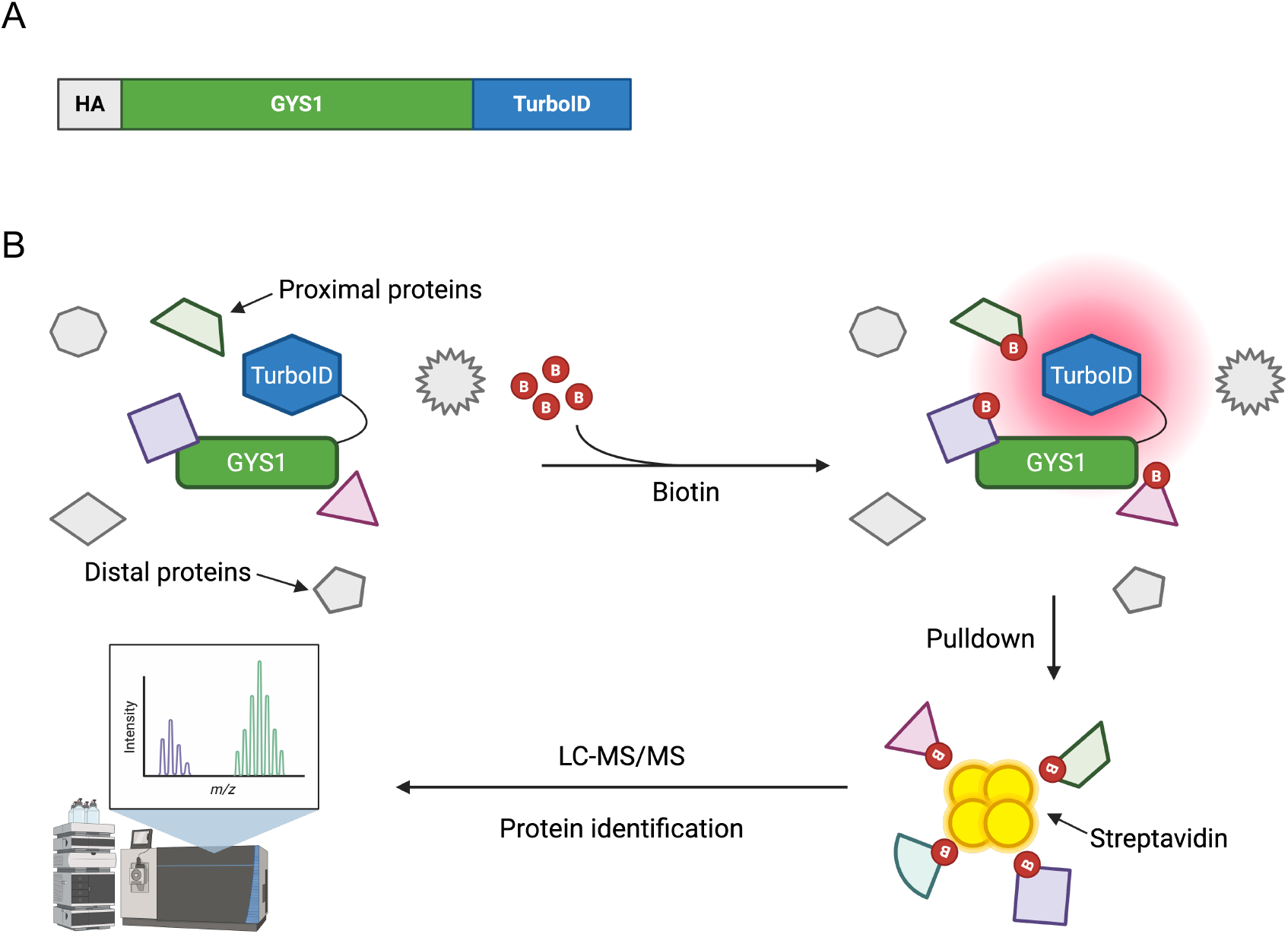
Identification of GYS1 interacting partners by TurboID-based proximity labeling. (A) In brown adipose tissue (BAT), glycogen metabolism is acutely regulated under metabolic shifts like thermogenic activation. Glycogen synthase 1 (GYS1), the rate-limiting enzyme of glycogen synthesis, may play a role in coordinating energy expenditure in brown adipocytes. (B) GYS1 TurboID fusion construct (GYS1-TbID). (C) Schematic overview of the TurboID-mediated proximity biotinylation assay using GYS1-TbID. The TurboID ligase uses biotin to covalently label neighboring proteins. Biotin-tagged proteins are isolated by co-immunoprecipitation with streptavidin-coated beads, then analyzed by LC-MS/MS for identification of GYS1 interacting partners. Figures created in BioRender.

## RESULTS

### GYS1-TbID is stably expressed in primary pre-BAT cells and exhibits functional biotinylation activity

To perform proximity labeling of GYS1 interacting partners by TbID, we generated two stable pre-BAT cell lines by MMLV retroviral transduction: one containing an empty pBABE vector (pBABE), and one containing an HA-tagged GYS1-TbID gene fusion construct (GYS1-TbID) (Figure 1A). Western blotting was conducted in pBABE and GYS1-TbID protein lysates to assess stable expression of the GYS1-TbID fusion construct in pre-BAT cells (Figure 2A). When probed for GYS1, endogenous GYS1 was detected in both cell lines at the expected ∼84kDa. Since TbID ligase carries a molecular weight of ∼35kDa, the presence of additional bands at ∼119kDa in GYS1-TbID cells only indicates expression of the fusion construct. To specifically detect expression of the fusion construct, lysates were probed for HA-tag. The HA epitope tag was detected in GYS1-TbID cells only at ∼120kDa, further confirming stable expression of the GYS1-TbID fusion construct.

**Figure 2.**
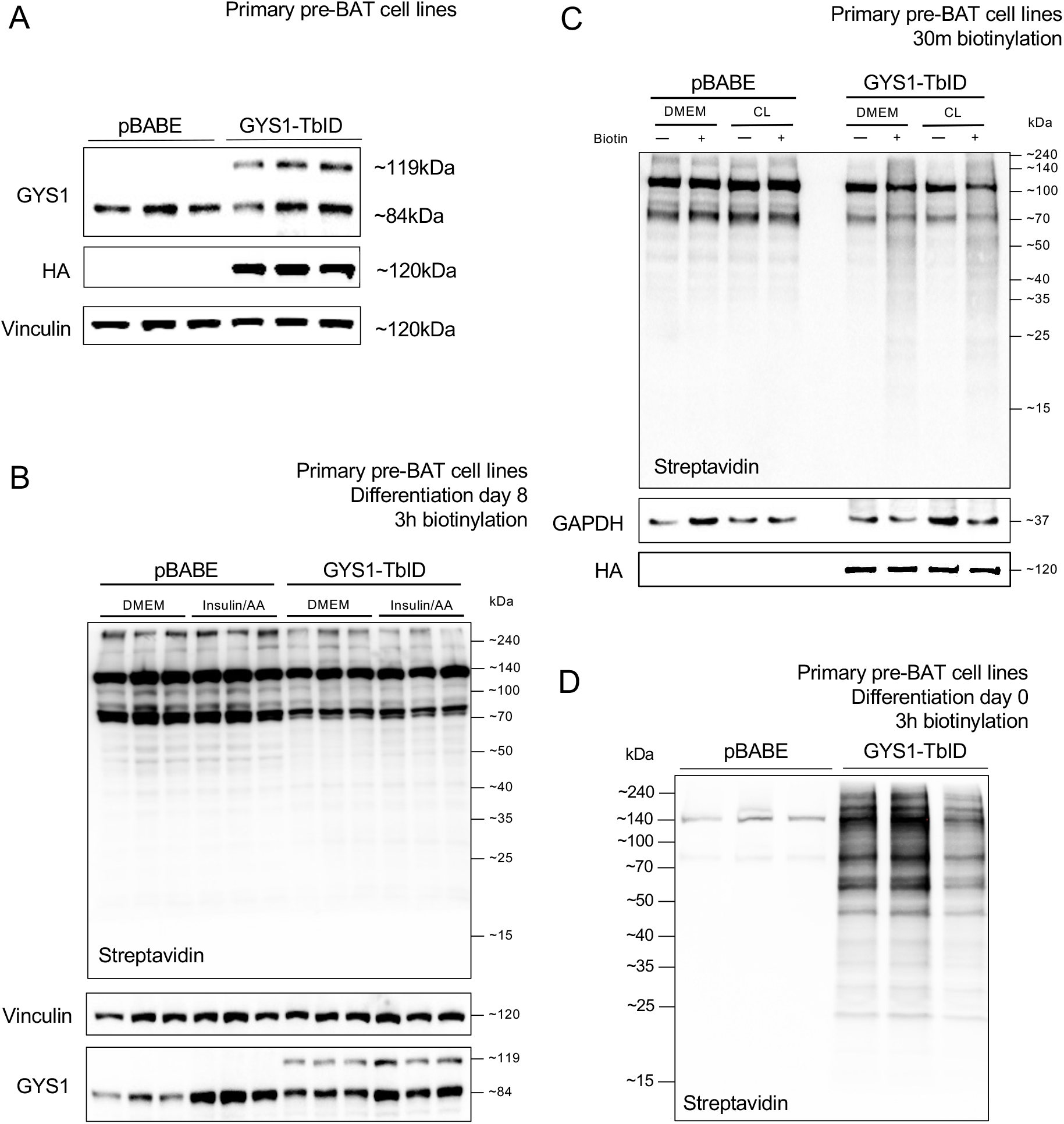
GYS1-TbID is stably expressed in pre-BAT cells and exhibits functional proximity labeling. (A) Expression of the specified proteins in undifferentiated pBABE and GYS1-TbID pre-BAT cells. GYS1-TbID lysates exhibit additional bands when probed for GYS1 due to expression of the GYS1-TbID fusion construct (TbID molecular weight ∼35kDa). (B) Expression of the specified proteins in pBABE and GYS1-TbID pre-BAT cells after 8 days of differentiation and 3 hours of biotin treatment. (C-D) Expression of the specified proteins in undifferentiated pBABE and GYS1-TbID pre-BAT cells after (C) 30 minutes or (D) 3 hours of biotin treatment.

To verify that the fusion construct retained functional TbID proximity labeling activity, we treated both pBABE and GYS1-TbID cells with biotin and assessed biotinylated proteins by streptavidin blotting (Figure 2B-D). Mature brown adipocytes exhibited substantial endogenous biotinylation, resulting in background signal in pBABE cells (Figure 2B). We next performed the assay in undifferentiated pre-BAT cells, which resolved the background noise (Figure 2C-D). Administering biotin for three hours yielded stronger enrichment than 30 minutes, so we used this condition for the final proteomics experiment.

### GYS1-TbID expression does not disrupt brown adipocyte differentiation

To verify that the fusion construct does not interfere with brown adipocyte function, the morphology of pBABE and GYS1-TbID cells were observed throughout differentiation into brown adipocytes by bright-field microscopy (Figure 3). Both cell lines exhibit fibroblast-like morphologies prior to differentiation (day 0). pBABE and GYS1-TbID cells accumulate lipid droplets at a similar rate throughout differentiation and become fully differentiated by day 8, suggesting that GYS1-TbID expression does not disrupt brown adipocyte differentiation.

**Figure 3.**
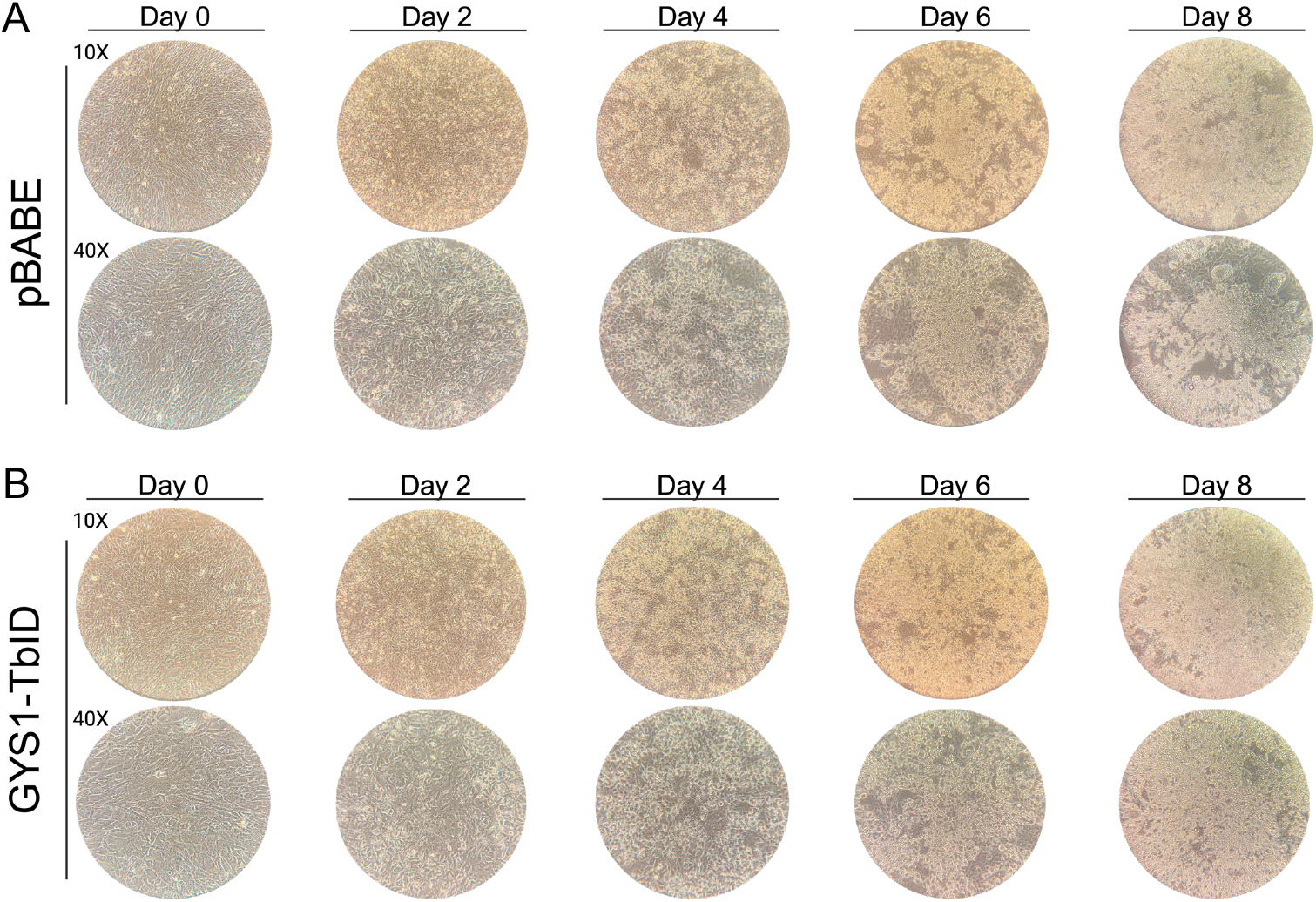
GYS1-TbID expression does not disrupt brown adipocyte differentiation. Representative bright-field microscopy images of (A) pBABE and (B) GYS1-TbID pre-BAT cells throughout differentiation into brown adipocytes.

### Proximal proteins labeled by GYS1-TbD are successfully enriched by co-immunoprecipitation

To enrich biotinylated proximal proteins labeled by GYS1-TbID for proteomic analysis, co-immunoprecipitation (coIP) was performed on whole-cell lysates using streptavidin-coated magnetic beads. Undifferentiated pBABE and GYS1-TbID pre-BAT cells were administered biotin for three hours before lysis. To optimize the bead-to-protein ratio for preparation of final proteomic samples, coIP was performed with increasing concentrations of streptavidin beads (10µL, 25µL, 50µL) with 300µg protein (Figure 4A). Streptavidin blotting with IP pulldowns revealed enrichment of biotin-tagged proteins from GYS1-TbID but not pBABE negative control samples. While higher streptavidin bead concentrations increase the yield of enriched material, they also amplify endogenous non-specific signals at 50µL. The most optimal bead-to-protein ratio for proteomic sample preparation was subsequently determined from these preliminary results to be 25µL per 300µg protein.

**Figure 4.**
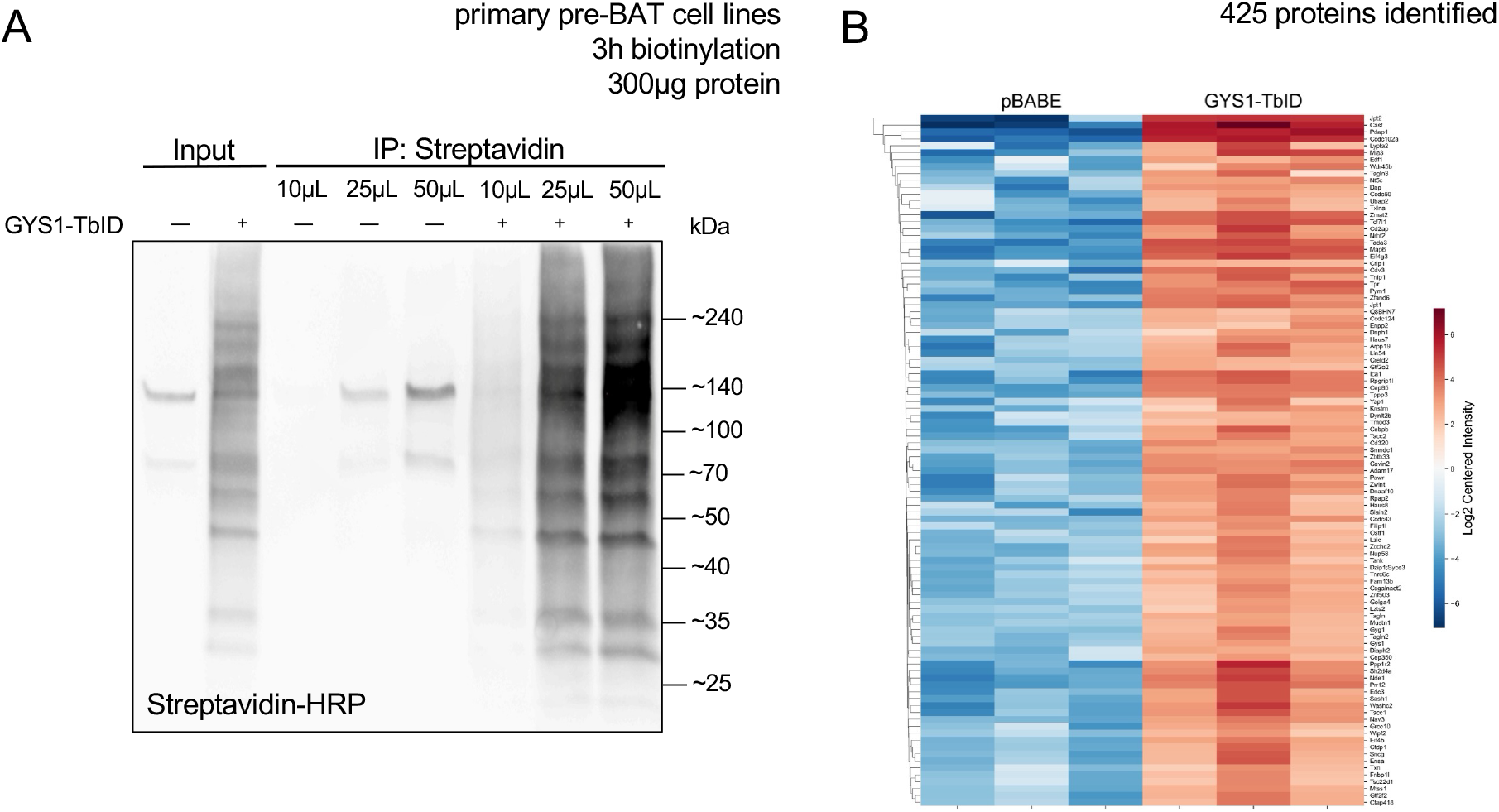
Enrichment and identification of biotin-tagged proteins by LC-MS/MS analysis. (A) Undifferentiated pBABE and GYS1-TbID pre-BAT cells were treated with biotin for 3 hours before lysis. Biotinylated proteins were enriched with increasing amounts of streptavidin beads (10μL, 25μL, 50μL) and detected by streptavidin blotting. (B) Hierarchical clustering of top 100 proteins significantly enriched in GYS1-TbID samples as determined by proteomic analysis (n=3 for each group).

### GYS1 interacting partners are enriched in multiple functions

For characterization of GYS1 interacting partners, IP samples were prepared following the above procedure in replicates and processed through mass spectrometry. Proteomic analysis identified 425 enriched proteins in GYS1-TbID cells (Figure 4B).

Among the identified interacting partners, many proteins were established glycogen-handling proteins or regulators of glycogen synthase activity (Figure 5A). Several of these proteins have been found to directly interact with GYS1, including GYG1, STBD1, and PP1 (24–26). Some identified glycogen-handling proteins, including WDR45B and PGM1, have not yet been characterized as GYS1 interacting partners and merit further experimental validation. In addition to known glycogen-associated proteins, some of the topmost enriched proteins included CAST, CCDC102A, TCF7L1, JPT1, and EIF4G3 (Figure 5B). Functionally, these proteins are involved in various cellular processes, including cytoskeletal dynamics, transcriptional regulation, intracellular signaling, and protein translation (27–31).

**Figure 5.**
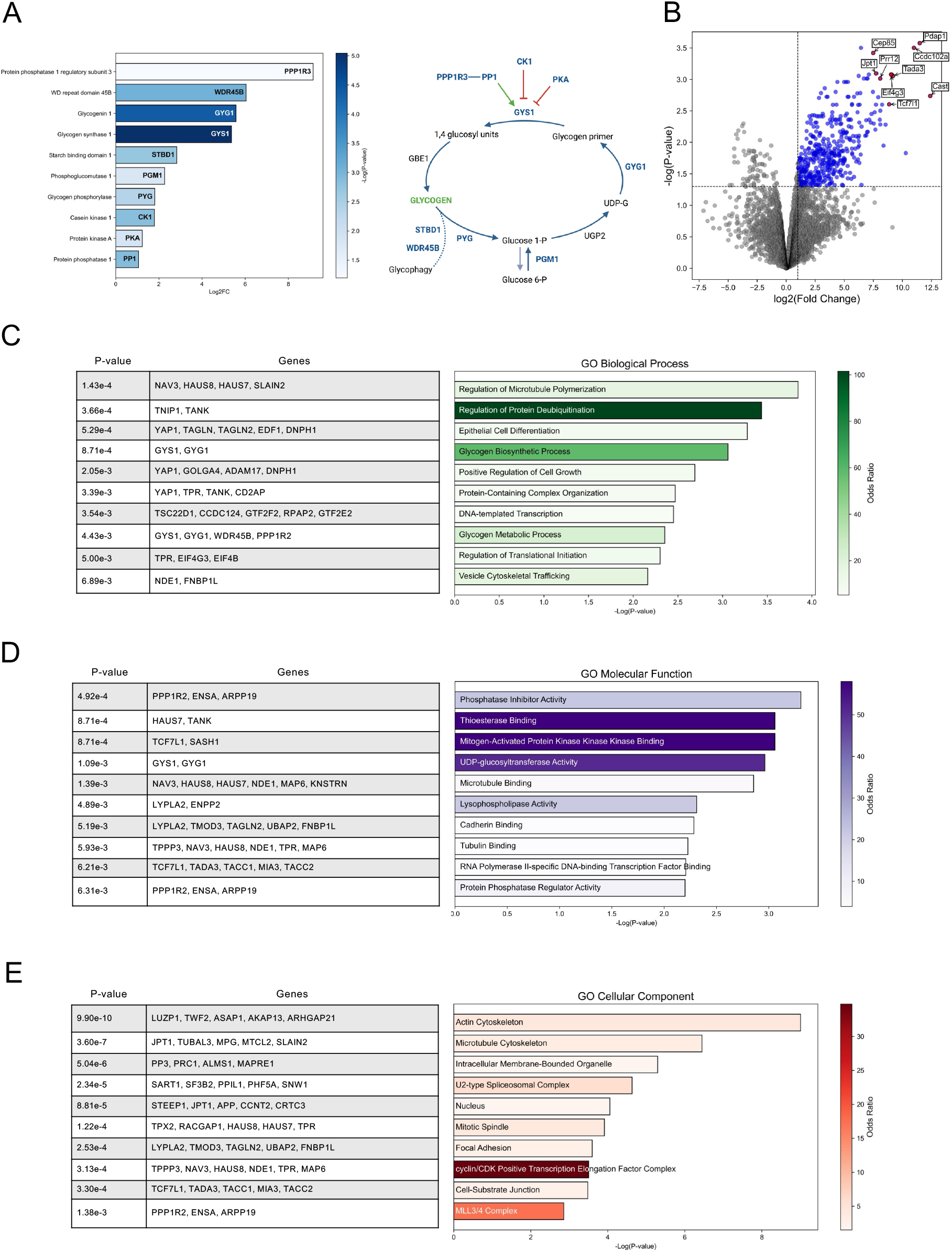
Identification of novel GYS1 interacting partners. (A) Volcano plot depicting proteins enriched in GYS1-TbID samples compared to pBABE negative control. Selected novel GYS1 interacting partners are labeled. (B) Glycogen metabolism enzymes and established GYS1 regulators identified by proximity labeling in GYS1-TbID samples. (C-D) Gene ontology enrichment analysis of the top 100 most enriched proteins for (C) biological process (D) and molecular function. (E) Gene ontology enrichment analysis of enriched proteins for cellular component. Analysis performed by Enrichr.

To further characterize the functions of the identified interacting partners, we performed pathway analysis on enriched proteins for biological process, molecular function, and cellular component through Gene Ontology (Figure 5C-E). In addition to enrichment for glycogen metabolism, GYS1 interacting partners are interestingly enriched in additional processes, including cytoskeletal remodeling, transcriptional regulation, and MAPK signaling. Interacting partners display surprising subcellular localization to the actin and microtubule cytoskeleton and the nucleus. Together, these results demonstrate that GYS1-TbID reliably labels proximal proteins in pre-BAT cells and reveal novel interacting partners involved in functions beyond glycogen metabolism.

## DISCUSSION

Glycogen is one of the primary mechanisms by which cells store and utilize energy (12). While its function is well-characterized in liver and skeletal muscle, our current understanding of its role in BAT remains limited. Characterizing the regulation of GYS1 in brown adipocytes can therefore help to illustrate the interplay between glycogen metabolism and BAT function. The TurboID system is optimized for efficient and non-toxic proximity labeling, rendering it most suitable for studying dynamic interactions with GYS1 (23). Our proteomic analysis of GYS1-TbID in pre-BAT cells reveals a protein-protein interaction network enriched not only for glycogen-handling proteins, but also for transcriptional regulators and cytoskeletal components. These findings suggest broader cellular functions for GYS1 beyond cytoplasmic glycogenesis.

Gene Ontology analysis of GYS1 interacting partners indicated unexpected enrichment in transcriptional regulators and nuclear localization. Several of these proteins were transcription factors, suggesting a role for GYS1 in gene regulation. This aligns with previous reports that GYS1 translocates between the cytosol and the nucleus in response to metabolic cues, as well as reports of its RNA binding capabilities (17,21). In particular, the transcription factors FoxO1, FRA2, TCF7L1, and YAP1 emerged as likely interacting partners, each with roles in regulating glucose or lipid metabolism. FoxO1 and FRA2 are central to metabolic homeostasis and adipogenesis, while TCF7L1 and YAP1 are downstream effectors of Wnt and Hippo signaling pathways (32–35). Further experimentation is required to validate these interactions with GYS1 and assess their subcellular localization.

In addition to transcriptional proteins, our data also indicate enrichment of proteins associated with cytoskeletal remodeling and localized to the actin cytoskeleton. Consistent with these findings, glycogen synthesis and resynthesis has been shown to be closely associated with actin filaments through the binding of glycogenin, which facilitates the formation of glycogen granules (36). These interactions may facilitate the spatial organization of intracellular glycogen during initial synthesis. These results, however, are correlative and warrant further investigation to determine their functional relevance.

Among the top candidates was the transcription factor Forkhead box O1 (FoxO1), a central downstream target of insulin signaling. Activation of the PI3K/AKT signaling pathway by insulin leads to the phosphorylation and nuclear exclusion of FoxO1, thereby suppressing its transcriptional activity (32). FoxO1 acts as a transcriptional modulator of genes involved in insulin sensing, lipid metabolism, mitochondrial activity, and energy uptake. It is abundantly expressed in the pancreas, liver, skeletal muscle, white and brown adipose tissue, and the hypothalamus. Functionally, FoxO1 promotes gluconeogenesis in the liver and lipid oxidation in the skeletal muscle during fasting, while in adipose tissue it suppresses adipogenesis (37–39). Notably, selective inhibition of FoxO1 in BAT enhances oxygen consumption and thermogenic gene expression, and hepatocyte-specific FoxO1 knockout decreases blood glucose while increasing glycogen content (40,41). The regulation of FoxO1 by AKT signaling, along with its abundance in adipose tissue and relevance to glycogen metabolism, supports a potential interaction between GYS1 and FoxO1. One intriguing possibility is that GYS1 may function to sequester FoxO1 in the cytosol under fed conditions, thereby preventing transcription of gluconeogenic genes.

To further characterize the association of GYS1 with candidate interacting partners like FOXO1, reciprocal co-immunoprecipitation (coIP) experiments should be performed to confirm endogenous protein-protein interactions. These experiments should also be performed under varying metabolic states and stages of differentiation to assess their effects on the strength and stability of GYS1 interactions. Loss-of-function studies using CRISPR/Cas9-mediated knockout of candidate partners will help to functionally characterize their roles in glycogen regulation. These findings will provide deeper insight into glycogen regulation and its role in brown adipocyte activation and metabolism.

## MATERIALS AND METHODS

### Cloning of GYS1–TurboID fusion

The coding sequence of GYS1 and TurboID were obtained from validated plasmid templates. Coding regions were amplified with a high-fidelity polymerase using primers designed to (i) remove the GYS1 stop codon, (ii) introduce a short flexible linker at the GYS1–TurboID junction, and (iii) add either homologous overlaps for Gibson-style assembly or att sites for Gateway recombination. For Gibson-style assembly, the expression vector was linearized to expose ends complementary to the insert overlaps, and assembly was performed according to the manufacturer’s recommendations. For Gateway cloning, the attB-flanked GYS1–TurboID insert was recombined into a donor vector via the BP reaction to generate an Entry clone, followed by LR recombination into destination vectors for expression. Following transformation into a standard cloning strain, colonies were screened by junction PCR and verified by Sanger sequencing across the entire fusion open reading frame and all junctions. Final constructs were archived as annotated sequence files and used for downstream expression tests.

### Generation of pre-BAT cell lines

Pre-BAT cells were isolated and immortalized as previously described (42). Stable pre-BAT cell lines expressing either an empty pBABE vector or the HA-GYS1-TbID fusion construct were generated via MMLV-based retroviral transduction.

### Cell culture

Cells were cultured in Dulbecco’s modified Eagle’s medium (DMEM) supplemented with 10% fetal bovine serum (FBS). For differentiation into brown adipocytes, pre-BAT cells were cultured to 100% confluency. Confluent pre-BAT cells were induced to brown adipocytes using adipogenic cocktail containing DMEM, 10% FBS, 0.5µM dexamethasone, 0.5mM IBMX, 0.125mM indomethacin, 2nM T3, 20nM insulin, and 1µM rosiglitazone. The medium was changed every 2 days with fresh media containing DMEM, FBS, T3, insulin, and rosiglitazone.

All cell culture experiments were performed in triplicate in 10cm dishes (1x10^6^ cells) or 6-well plates (2x10^5^ cells). For biotin labeling, confluent pre-BAT cells were starved for 1 hour in 1x HBSS, then administered 500µM biotin for 3 hours before harvesting.

### Protein lysis

Total protein was extracted in Radio-Immunoprecipitation Assay (RIPA) lysis buffer supplemented with protease/phosphatase inhibitor cocktail. For protein lysis following biotin labeling, cells were washed 5 times in ice cold 1x PBS before lysis with RIPA buffer. Lysates were incubated at 4°C for 20 minutes, then transferred into 2mL Eppendorf tubes and vortexed twice. Samples were ultrasonicated twice for 20 seconds at 10% amplitude, then centrifuged at 17,000g for 20 minutes at 4°C. The supernatant was collected and stored at -80°C until analysis.

### Western blotting

Protein concentrations were determined by the Pierce™ BCA Protein Assay Kit or by Bradford protein assay. 20µg of total protein extract were separated by 10% SDS-PAGE, then transferred to nitrocellulose membranes at 100V for 1 hour. Membranes were blocked in 5% milk blocking buffer or 5% bovine serum albumin (BSA) for 1 hour at room temperature, then incubated with primary antibodies (1:1000) at 4°C overnight. Membranes were incubated with horseradish peroxidase-conjugated secondary antibodies (1:5000) for 1 hour at room temperature the next day, and Western blots were developed using Azure Radiance Plus Chemiluminescent substrate and detected by Azure Sapphire Biomolecular Imager. For streptavidin blot analysis, membranes were incubated with streptavidin-HRP antibody (1:3333) for 30 minutes at room temperature before developing.

Glycogen synthase 1 and streptavidin-HRP antibodies were purchased from Cell Signaling Technologies. Monoclonal anti-vinculin antibody was purchased from Millipore Sigma.

### Co-immunoprecipitation

For IP-MS optimization, whole-cell lysates (300µg protein per sample) were incubated with streptavidin-conjugated magnetic beads at 4°C overnight. Beads were washed with RIPA, 1M KCl, 0.1M Na2CO3, and 2M urea in 10mM Tris-HCL (pH 8.0) at room temperature the following day. Enriched material was eluted from the beads by boiling in 3x SDS loading buffer supplemented with 2 mM biotin and 20 mM DTT at 95°C for 10 minutes. Streptavidin-HRP blotting and total-protein imaging (using stain-free gels) were performed on input and pulldown samples to verify enrichment.

For preparation of proteomic samples for mass spectrometry, coIP was performed on biotin-labeled lysates following standard protocol, without elution. Bound beads were resuspended in 2M urea in 100mM Tris-HCL (pH 8.0) for on-bead digestion and mass spectrometric analysis.

Protein-bound beads were resuspended in 2 M urea containing 50 mM Tris-HCl (pH 8.0), followed by reduction with 5 mM tris(2-carboxyethyl)phosphine (TCEP) and alkylation with 10 mM iodoacetamide. Proteins were digested overnight at 37 °C with lys-C and trypsin. The digestion reaction was quenched by adding formic acid to a final concentration of 5% (v/v). Peptides were cleaned using the SP3 protocol (43), dried, and reconstituted in 5% formic acid for LC-MS/MS analysis.

### Mass spectrometry

Liquid chromatography–mass spectrometry was carried out on a Vanquish Neo UHPLC system coupled to an Astral mass spectrometer (Thermo Fisher Scientific, Bremen, Germany). Samples were loaded and separated using a trap-and-elute workflow on a PepSep C18 column (150 mm × 150 µm, 1.7 µm particles) maintained at 59 °C. Mobile phase A was water with 0.1% formic acid, and mobile phase B was acetonitrile with 0.1% formic acid. Peptides were separated over a 15-minute gradient: 5% B from 0–1 min (2.45 µL/min), 5–15% B from 1–5 min (1.75 µL/min), 15–25% B from 5–12.6 min (1.75 µL/min), 25–38% B from 12.6–13.6 min (1.75 µL/min), 38– 80% B from 13.6–13.7 min (2.45 µL/min), and held at 80% B until 15 min (2.45 µL/min).

Data-independent acquisition (DIA) was performed on the Astral mass spectrometer in positive electrospray ionization mode. MS1 spectra were acquired at 240,000 resolution across m/z 380–980 with a normalized AGC target of 500% and a 3 ms maximum injection time. DIA windows were 4 m/z wide across the 380–980 m/z range, with MS2 spectra acquired at 80,000 resolution using 25% normalized HCD, a normalized AGC target of 500%, and a 7 ms maximum injection time. Raw data were processed with DIA-NN using an in-silico spectral library generated from a UniProt mouse proteome database (44). DIA-NN results were further analyzed in FragPipe Analyst to identify differentially enriched proteins (45). Data have been deposited to the MassIVE repository under accession number MSXXXXXX.

### Pathway analysis

To identify significantly enriched Gene Ontology (GO) terms of enriched interacting partners, we used Enrichr, an integrative web-based application (46–48). Enriched GO terms with adjusted p-value < 0.05 and involving at least 2 genes were considered statistically significant.

